# Developmental stages of *Arthroleptella villiersi* (Pyxicephalidae)

**DOI:** 10.1101/2025.03.31.646409

**Authors:** Susan Schweiger, Benjamin Naumann, Hendrik Müller

## Abstract

Terrestrialization in anurans is associated with the evolution of endotrophy. It is hypothesized that heterochrony, or changes in the time or rate of developmental events, is associated with the development in endotrophic species. To analyse the developmental heterochrony, we investigated and revised the description of the development in *Arthroleptella villierisi*, a small frog species of the family Pyxicephalidae, found in the Cape fold mountain region of the Western Cape, South Africa. We also compared the developmental stages of *A. villiersi* to the heterochronic patterns of taxa with endotrophic terrestrial indirect and direct development using heterochrony plots to identify heterochronic shifts during development. As a result, we found that the terrestrial endotrophic larva of *A. villiersi* shares external similarities with exotrophic, aquatic larvae in having a long muscularized tail with a fin, a lateral line system and an opercular fold that completely covers the forelimbs. However, other developmental events like the reduction of larval mouthparts and the pre-displaced fore- and hindlimb development is comparatively similar to direct developing taxa. The results of our study show that the timing of early developmental events can be shifted profoundly, while the timing of later events seem to be more conserved in anuran development. We interpret that some of these heterochronic shifts might be consequences of functional and developmental constraints underlying the establishment of the adult body plan.

## Introduction

All extant anurans originate from a last common ancestor with an indirect-aquatic reproductive mode including a free-living and feeding (exotrophic) larva called tadpole (Chuliver et al., 2024). These larvae actively feed and exhibit a wide range of morphological and behavioural adaptations depending on their habitat preferences (Altig & Johnston, 1989; Wells 2007). The aquatic tadpole undergoes a climactic process of morphological and physiological rearrangement (metamorphosis) to develop into a more or less terrestrial adult (Duellman & Trueb, 1994). This ancestral mode has been modified in several anuran lineages towards a partial or complete terrestrialization of reproduction, resulting in various degrees of independence from aquatic habitats (e.g. Noble 1931; Haddad & Prado 2005; Crump 2015). This shift towards terrestrialization was facilitated by the evolution of endotrophy, in which embryos and tadpoles rely on their yolk reserves rather than on obtaining nutrients from the external environment (Thibaudeau & Altig 1999). Endotrophy is found in anurans with different reproductive modes ranging from indirect development with a free-living, non-feeding (“nidicolous”) larva to direct development, which is completed inside a terrestrially laid egg and results in the hatching of a fully formed froglet, thus bypassing the free-living aquatic larval stage (Callery & Elinson 2001, Thibaudeau & Altig 1999, Altig & Johnston 1989). In lineages with the complete terrestrialization of reproduction, endotrophy is regarded as a key innovation allowing the large-scale repatterning of embryonic and larval development (Elinson, 2009; Hanken, 2006). This repatterning can be described in several ways, including changes in space (heterotypy), amount (heterometry) and time (heterochrony). Among these changes, heterochrony, the phylogenetic differences in developmental rate and timing, has been hypothesized to be exceedingly important (Grosso et al., 2019; Vera Candioti et al. 2011; Hanken et al. 1992, Raff & Wray 1989).

However, comparative qualitative analyses following a formalized method to test this hypothesis have not been carried out. Detailed developmental staging tables for terrestrial endotrophic anurans are crucial to for the analyses of developmental heterochrony. They are the basis for a formalized comparison of the timing of homologous developmental events and allow the identification of heterochronic shifts. However, both, the generalized staging system for exotrophic tadpoles, established by Gosner (1960), and the system of Townsend & Stewart (1985) established for *Eleutherodactylus* and often applied to endotrophic direct developing taxa, are unsuitable for many endotrophic indirect developing species. These forms are characterized by terrestrial deposition of eggs and a free-living larva that hatches from the egg but remains within a terrestrial nest, where it does not feed until metamorphosis is completed (Wells 2007). This mode of development has evolved independently in several taxa (Thibaudeau & Altig 1999). The morphological modifications of such endotrophic larvae range from tadpoles that are superficially similar to free-living, exotrophic tadpoles, to highly modified terrestrial larvae that lack many of the tadpole-typical features (Juncá et al. 1994, Vera Candioti et al. 2011).

Such a form of indirect development with terrestrial, endotrophic tadpoles is found in species of *Arthroleptella*, a genus of small frogs in the family Pyxicephalidae (de Villiers 1929, van der Meijden et al. 2011) (Figure 1A). *Arthroleptella* currently comprises ten species (Turner & Channing, 2017), which are all confined to montane fynbos habitats of the Cape fold mountain region of the Western Cape, South Africa (Channing 2004a). The most common of these, De Villiers’ Moss Frog, *Arthroleptella villiersi* Hewitt, 1935, is found in the Hottentots Holland, Kogelberg, Kleinrivier Mountains and east to the Bredasdorp Mountains, where they are locally abundant and considered to be of Least Concern from a conservation perspective (IUCN SSC Amphibian Specialist Group, 2013). *Arthroleptella villiersi* live and breed in the fynbos vegetation in the foreground and lays small clutches of about 11 eggs in secluded places under moss or other vegetation, or hidden within loose soil or other organic matter, where they will complete development within the terrestrial nest (Channing 2004b) (Figure 1 B-D). Since its first description by de Villiers (1929), this highly derived form of development has been little studied except for Morgan et al.’s (1989) study of the endocrine mechanisms of metamorphosis in *A. lightfooti*.

**Figure 1.**
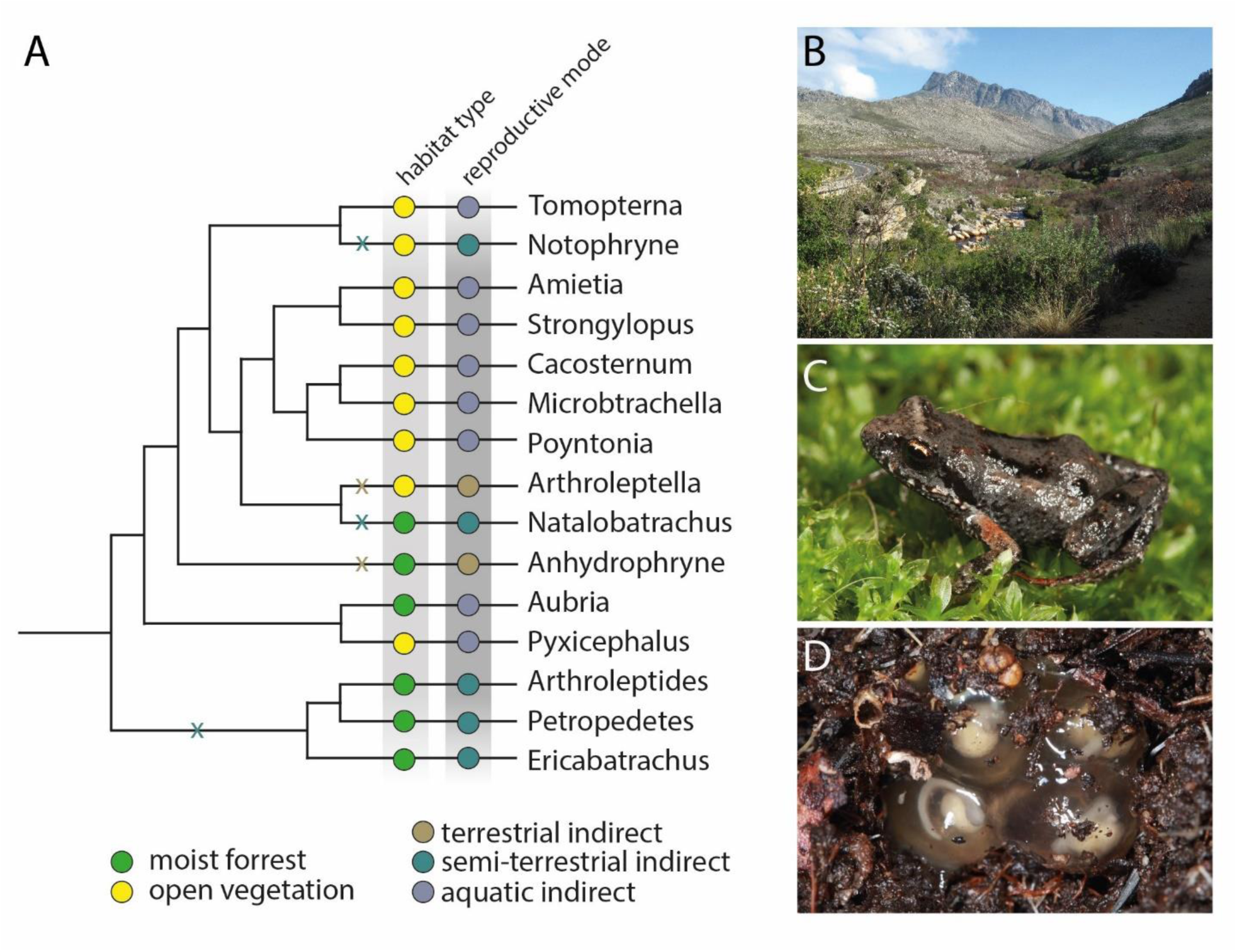
(A) Reconstruction of the phylogeny, habitat types and breeding modes of Pyxicephalidae. Modified from Bittencourt-Silva et al. 2016. (B) Typical habitat of *Arthroleptella villiersi* at Jonkershoek Nature Reserve, where the frogs live and breed in the fynbos vegetation in the foreground. (C) Adult *A. villiersi.* (D) Terrestrial clutch of *A. villiersi.* Photographs by Hendrik Müller.

We here revise the description of the developmental stages in *A. villiersi* and provide stage diagnostic characters for a comprehensive description of developmental stages. Additionally, we comment on the distinction of indirect terrestrial development and direct development and compare the heterochronic patterns of taxa with endotrophic terrestrial indirect and direct development using heterochrony plots (Schlosser and Roth, 1997, Schlosser 2001).

## Materials & Methods

### Specimens

A total of 95 embryos and hatchlings and two adults of *Arthroleptella villiersi* were collected between 2011 and 2014 near Stellenbosch, Kleinmond, and Jan Joubertsgat Bridge, Western Cape Province, South Africa. We transferred clutches to petri dishes and maintained them under field conditions at ambient temperatures, sampling embryos at regular intervals (every 1-2 day). Specimens were euthanized using tricaine methanesulfonate (MS222; Fluka), fixed in 4% buffered formalin, and subsequently stored in 70% ethanol. Conspecificity of the egg clutches was confirmed through identification of the attending male (du Preez and Carruthers, 2017) and by raising individual eggs to hatching. Descriptions of embryo morphology and their external features are based on microscopic observations of preserved material. Photos were made using a Zeiss Discovery V12 stereo microscope (Carl Zeiss) with an attached drawing mirror and an AxioCam digital camera with associated imaging software (Axio Vision).

### Heterochrony plot

In a heterochrony plot, the timing of developmental events in one species on the x-axis is plotted against the timing of development of homologous events in another species on the y-axis (Schlosser, 2001; Schlosser & Roth, 1997). This graphical method allows the detection of conserved timings of events as well as altered timings (pre-displacement and post-displacement). We conceptualized 34 developmental events and determined their developmental timing from staging systems available in the literature or, for *A. villiersi*, in the present study. For a detailed list of the events and the corresponding developmental timing in the different anuran species investigated in this study as well as the literature used to determine them see Table 1. Events assumed to be causally linked were grouped into eight suites indicated by roman numbers I to VIII (Table 1). Events for which the timing was not available for a species examined were excluded from the plot between two species. Regression lines are given for the suites to estimate developmental rates. The closer all events of a developmental suite fit to a linear regression, the more similar are the developmental rates of these events between the two species. Error bars were determined for each suite as proposed by Schlosser, 2001. To facilitate a more comprehensive comparison, four corresponding ontogenetic phases were defined based on very general developmental characters and indicated by dotted lines in each plot. Phase 1 (“embryonic” phase) covers the period from fertilization until before the emergence of optic and gill arch bulges. Phase 2 (“extended pharyngula” phase) the first appearance of optic and gill arch bulges as well as the elongation of the tail bud until before the visibility of the pupil, the darkening of the iris as well as the maximum extension of the operculum (additionally the presence of forelimb buds beneath the operculum and a paddle-shaped foot in direct-developing species). Phase 3 (“tadpole” phase) reaches from the presence of the characters until before first signs of tail regression. Phase 4 (“metamorphosis” phase) spans the onset of tail regression until one stage before the completion of metamorphosis defined by the complete disappearance of a tail stub. For the exact assignment of developmental stages of each species compared within the plots see Table 2.

**Table 1:**
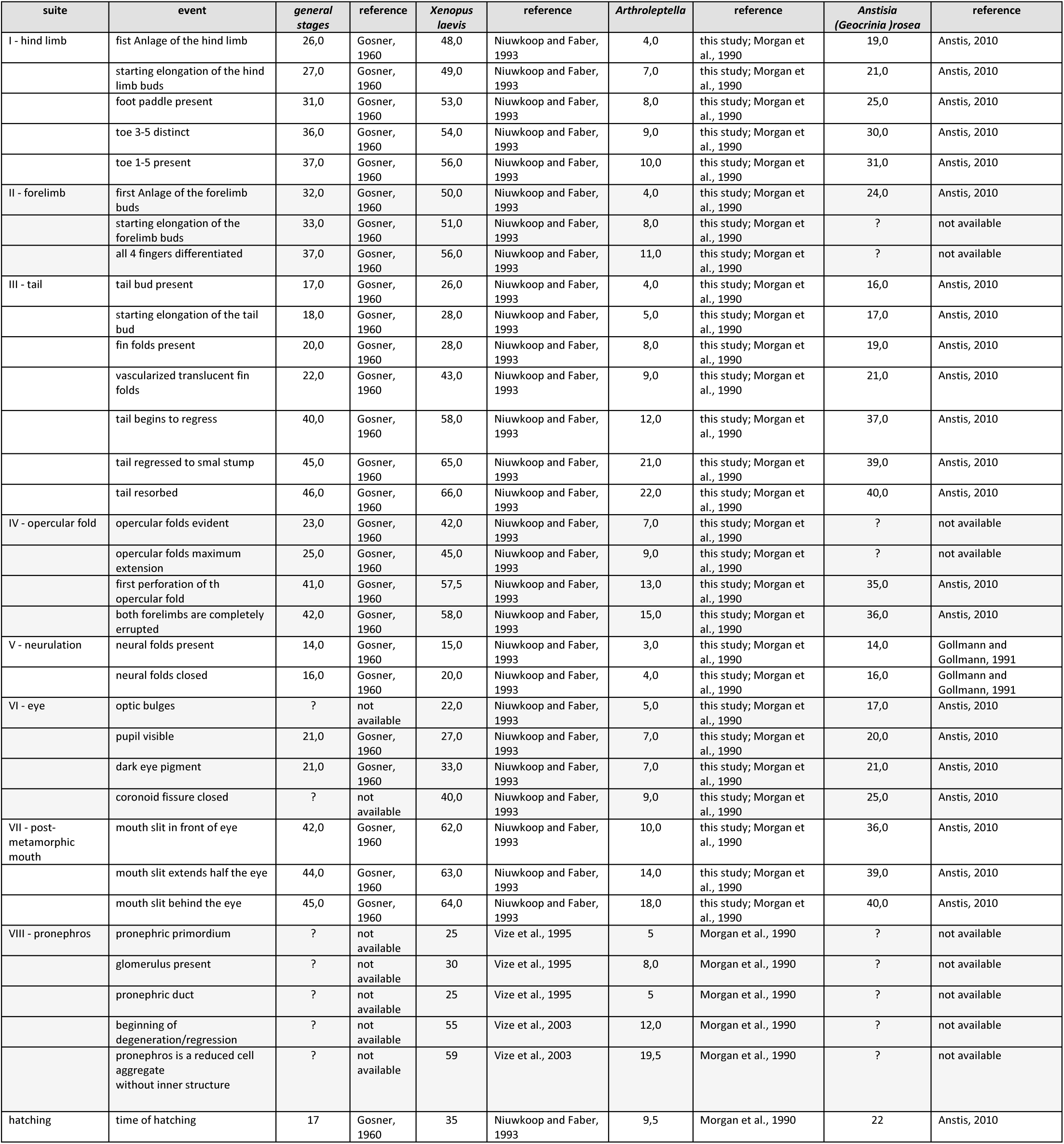
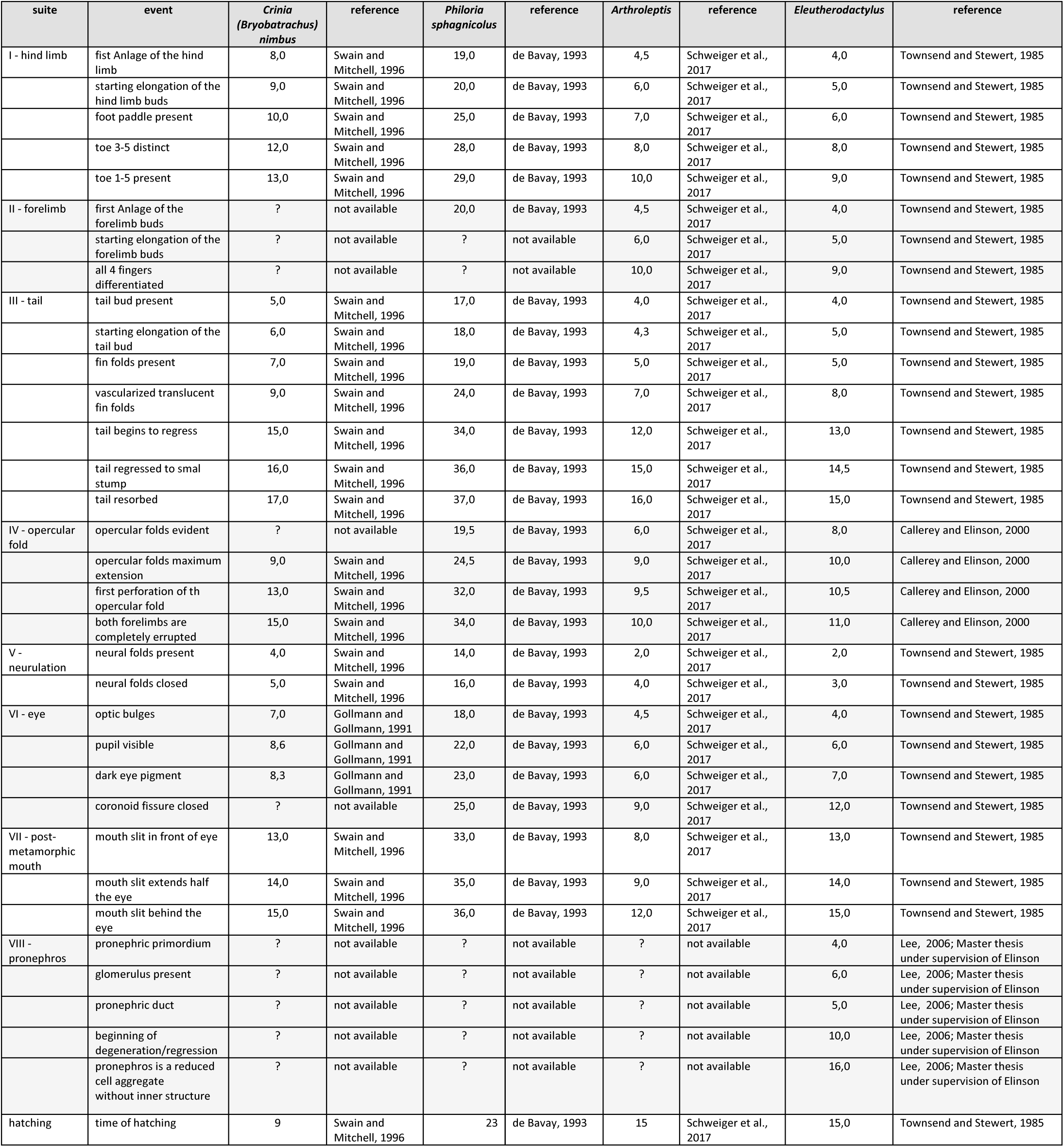
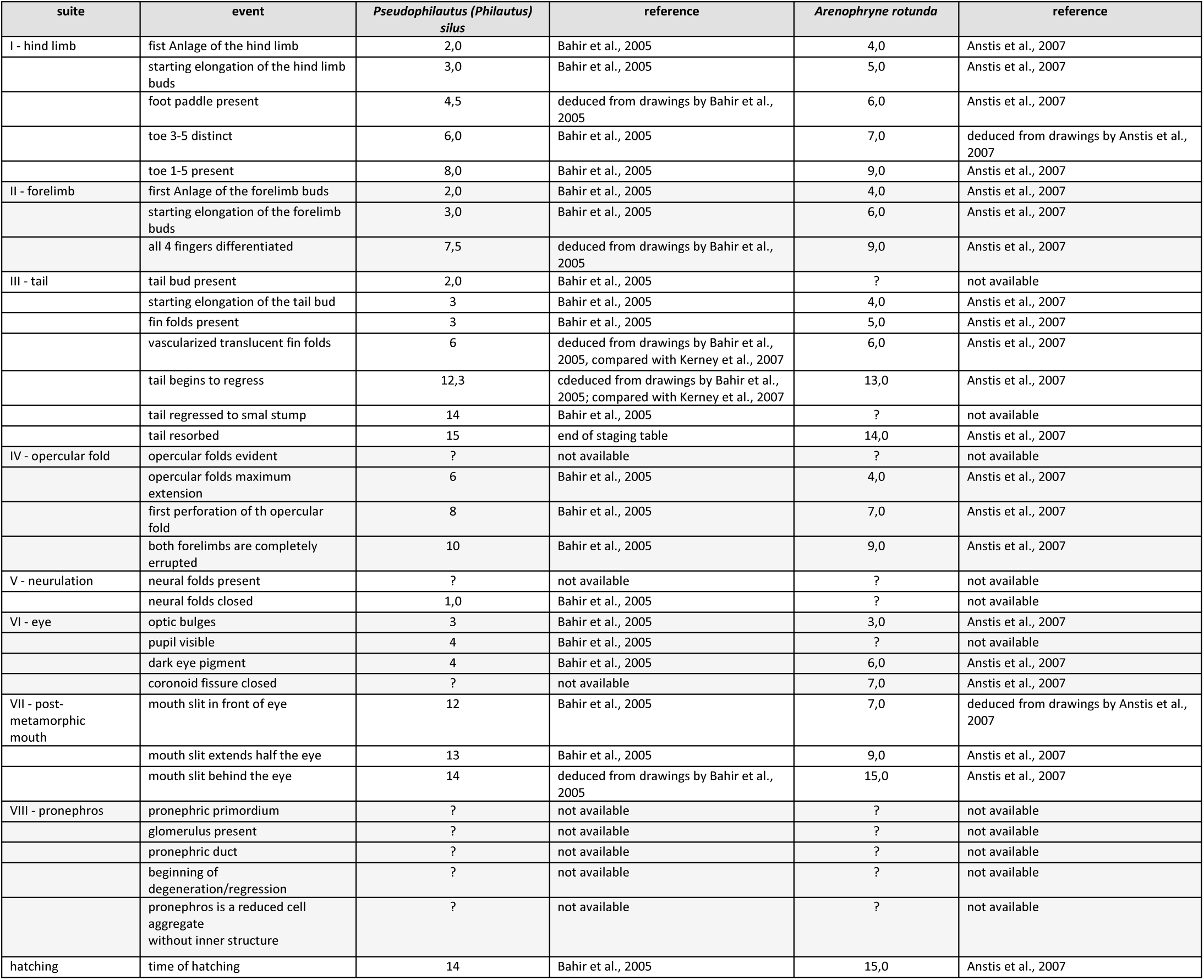
Developmental events of different anuran species compared in the heterochrony plots.

**Table 2:** Ontogenetic phases compared in the heterochrony plots.

| ontogenetic phase | *Arthroleptella*, present study | *Arthroleptella* Morgan et al., 1990 | *Arthroleptella* Gosner, 1960 (by Morgan et al., 1990) | comparative characters (this study) | Gosner, 1960 (this study) | *G. rosea* (Anstis, 2010) | *B. sphagnicola* (de Bavay, 1993) | *B. nimbus* (Swain and Mitchell, 1996/de Bavay-stages) | *E. coqui* (Townsend and Stewart 1985) | *A. wahlbergii* (schweiger et al., 2017) |
| --- | --- | --- | --- | --- | --- | --- | --- | --- | --- | --- |
| "embryonic"-phase | 3 | - | 15-16 |  |  |  |  |  |  |  |
|  | 4 | 1 | 27-28 |  |  |  |  |  |  |  |
|  | 5 | - | -(29?) |  |  |  |  |  |  |  |
| "hatchling"-phase | 6 | - | -(29?) | optic bulge; gill arches present; tail elongated | 20 | 20 | 20 | 8/20 | 4 | 4 |
|  | 7 | - | 29? |  |  |  |  |  |  |  |
| "tadpole" phase | 8 | 1.5 | 30 | pupil visible, iris dark/black; forelimb buds under opercular fold (maximum extension of the operculum); (foot paddle-shaped) | 25 | 25 | 25 | 10/25 | 7 | 7 |
|  | - | 2 | 31 |  |  |  |  |  |  |  |
|  | 9 | 2.5 | 32-33 |  |  |  |  |  |  |  |
|  | 9-10 | 3 | 34 |  |  |  |  |  |  |  |
|  | 10 | 3+ | 35-36 |  |  |  |  |  |  |  |
|  | - | 3.5 | 37 |  |  |  |  |  |  |  |
|  | 11 | 4- | -(38-40?) |  |  |  |  |  |  |  |
| "metamorphosis"-phase | 12 | 4 | -(38-40?) | begin of tail regression | 40 | 37 | 32-33 | 13-14/32-33 | 13 | 10-11 |
|  | 13 | 4+ | 41 |  |  |  |  |  |  |  |
|  | 14 | 4.5 | 42 forelimb; 43-44-adult mouth |  |  |  |  |  |  |  |
|  | 15 | 5- | 42 forelimb; 44-45-adult mouth |  |  |  |  |  |  |  |
|  | 16 | 5 | 45 |  |  |  |  |  |  |  |
|  | 17 | - | 45 |  |  |  |  |  |  |  |
|  | 18 | 5+ | 45-46 |  |  |  |  |  |  |  |
|  | 19 | 5.5 | 45-46 |  |  |  |  |  |  |  |
|  | - | 6- | 45-46 |  |  |  |  |  |  |  |
|  | - | 6 | 45-46 |  |  |  |  |  |  |  |
|  | 20 | 6.5 | 45-46 |  |  |  |  |  |  |  |
|  | 21 | 7 | 45-46 |  |  |  |  |  |  |  |
| post-metamorphosis-phase | 22 | - | 46 | completed metamorphosis | 46 | 40 | 37 | 17 | 15 | 16 |

## Results

### A revised staging system/table for species of *Arthroleptella*

Developmental stages of *A. lightfooti* have been loosely defined by Brink (1936) and later revised by Morgan et al. (1989). However, most of the characters used to define developmental stages focus on the progressive development of the tail, fore- and hind limbs, and developmental stages are not entirely depicted to facilitate further comparison with other taxa. In the present study, 20 developmental stages of the closely related *A. villiersi* are defined that describe overall morphological changes of embryos, hatchlings and larvae until the end of metamorphosis. The earliest developmental stages (from oviposition to the formation of the blastopore) have not been observed. A detailed description of the characters for each developmental stage is given in Table 3.

**Table 3:**
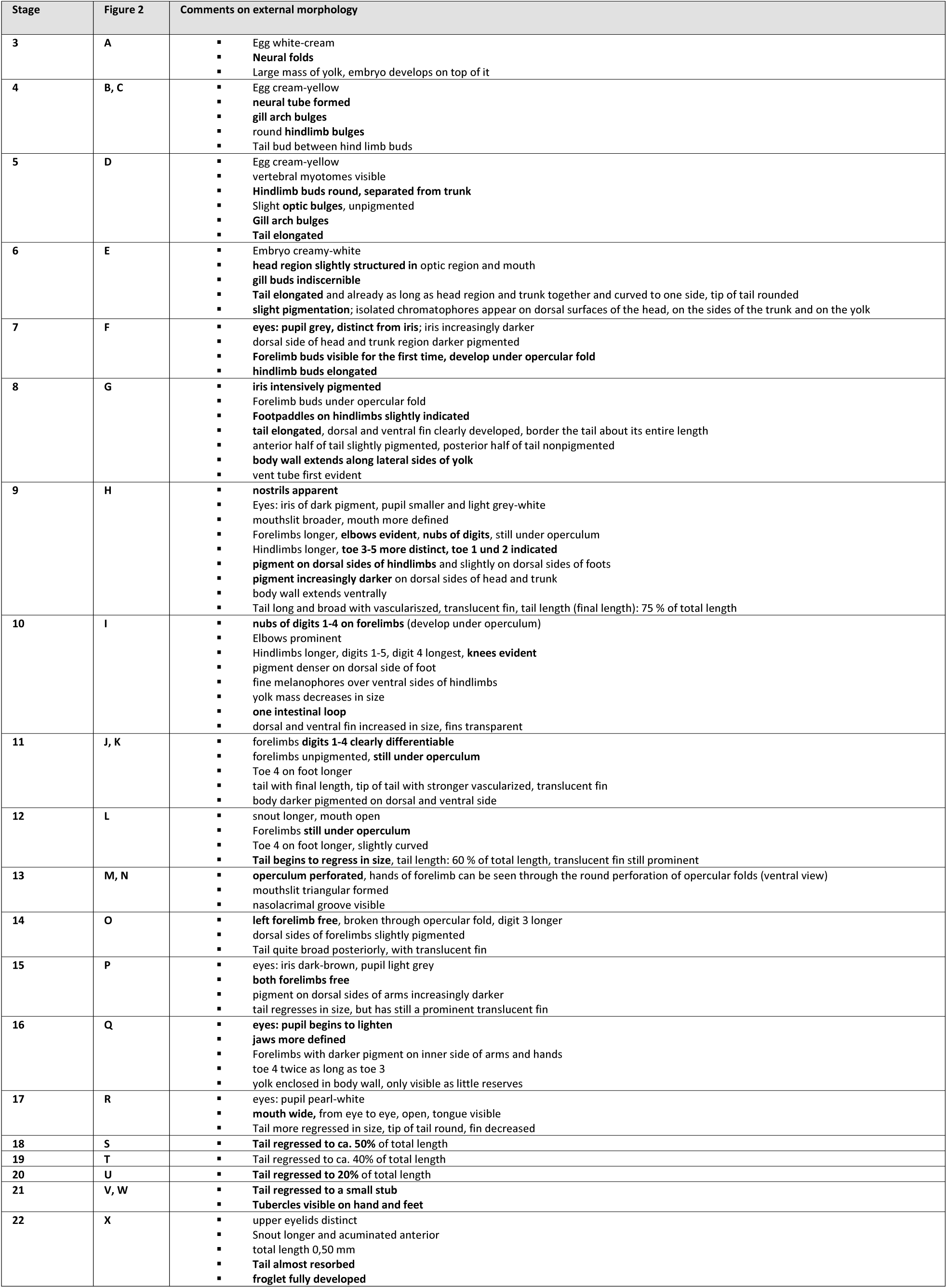
Staging table of *Arthroleptella villiersi*.

### General morphological development and metamorphosis

#### Head

The head is differentiated from the rest of the body at stage 4 (Figure 2B, C). Gill arch bulges are present till stage 5 (Figure 2D) and disappear by stage 6 (Figure 2E). No external gills were observed during development. Eye bulges became first visible at stage 5 but remain unpigmented during stage 6 (Figure 2D, E). At stage 7, the iris is indicated and becomes intensively pigmented at stage 8 (Figure 2F, G). Eyelids appear during stage 16 (Figure 2Q). Nostrils appear during stage 8. The mouth slit is indicated at stage 9. Upper and lower jaws become visible, and the mouth opens at stage 13 (Figure 2N). By stage 17, the mouth opening extends backwards below the eye (Figure 2R).

**Figure 2.**
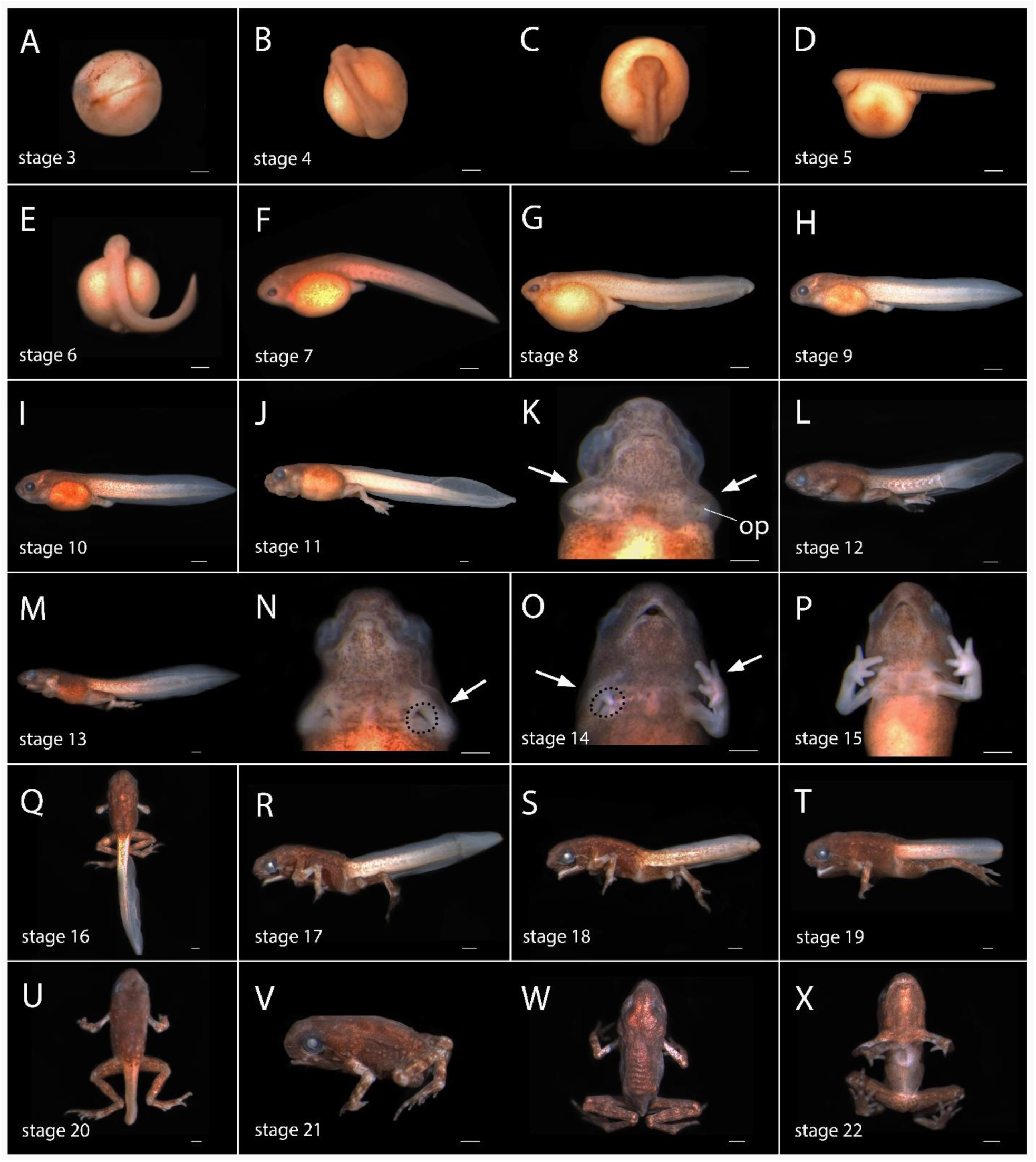
Stages of development in *Arthroleptella villiersi*. Embryos have been removed from the egg capsules. Arrows indicate developing forearms under an opercular fold (op). Scale bar 0.5 mm.

#### Body pigmentation and body wall

The eggs are creamy white to yellow. Light pigment appears along the trunk at stage 5 and begins to progress at stage 7, where melanophores are scattered all over the dorsal part of the head, trunk and 2/3 of the tail as well as the yolk (Figure 2D, F). By stage 9, pigmentation becomes increasingly darker and melanophores are widely spread over the ventral part of the head and the yolk and along the anterior half of the tail musculature (Figure 2H). The inner sides of the fore-and the hind limbs were lightly pigmented at stage 14 and increased in darkness during the following stages (Figure 2O-X). The upper and lower fin of the tail remain unpigmented during development.

#### Limbs

Both fore- and hind limb buds are present by stage 4, but the hind limb buds are more than twice as long as the forelimb buds (Figure 2B, C). The fore- and hind limb buds were joined to the trunk by stage 7 (Figure 2F). At this stage, the opercular folds are faintly indicated at the base of the forelimb buds and the hind limb buds are elongated. The forelimb buds are elongated and are completely covered by the opercular folds by stage 9 (Figure 2H). At the same stage, foot paddles of the hind limbs are first evident. At stage 10, elbow joints and knee joints are visible and nubs of toes 1-4 first appeared. The differentiation of hands and fingers was completed under the opercular folds during the following stages. By stage 13, a circular perforation of the opercular fold is initially visible on the ventral side (Figure 2M, N). By stage 14, the left forearm is fully erupted, and the circular perforation of the right forearm is visible (Figure 2O). By stage 15, both forearms completely erupted and fingers 1-4 are developed (Figure 2P). The subarticular tubercles of hands and feet are indicated at stage 20 and are clearly differentiated by stage 22 (Figure 2X).

#### Tail

The tail bud is first seen at stage 4 (Figure 2B). It has been elongated by stage 5 and bent to the right side of the body by stage 6 (Figure 2D, E). A fine fin has developed at stage 7 (Figure 2F). The tail has further elongated and reached its final length at stage 11 (Figure 2J). At this stage, it is well muscularized with a vascularized membranous fin. It begins to decrease during stage 15, when both forearms erupt through the opercular folds. At stage 20, the tail was greatly reduced and completely resorbed at stage 22 (Figure 2U-X).

### Heterochronic patterns underlying the evolution of endotrophic development in anurans

To compare patterns of heterochronic shifts underlying the evolution of endotrophic development, developmental events of different endotrophic species were plotted against homologous events in the exotrophic indirect developing *Xenopus laevis* (Figure 3A). First, we attempted to determine whether *X. laevis* can be used as a plesiomorphic representative of the outgroup. Therefore, developmental events of *X. laevis* were plotted against homologous events described in the generalized staging table for anurans (based on data from the bufonid *Incilius valliceps*) by Gosner (1960). Since the staging table by Gosner is simplified to fit the majority of exotrophic anurans it was only possible to determine the timing for some developmental events defined in the present study. The plot of *X. laevis* against the generalized Gosner-stages revealed the grouping of the tail, opercular fold, post-metamorphic mouth, forelimb and hind limb developmental suites into one isochronic suite B (Figure 3B, Table 4). Only three events exhibit clear telescoping (sequence heterochronic) shifts with two pre-displaced events (first perforation of the opercular fold, pupil visible) and one post-displaced event (hatching) identified in *X. laevis* compared to the generalized Gosner stages (Table 4). Based on this low number of heterochronic shifts compared to Gosner-stages and the more complete information on developmental events for *X. laevis* it is used as an exotrophic representative of the outgroup in subsequent plots.

**Figure 3.**
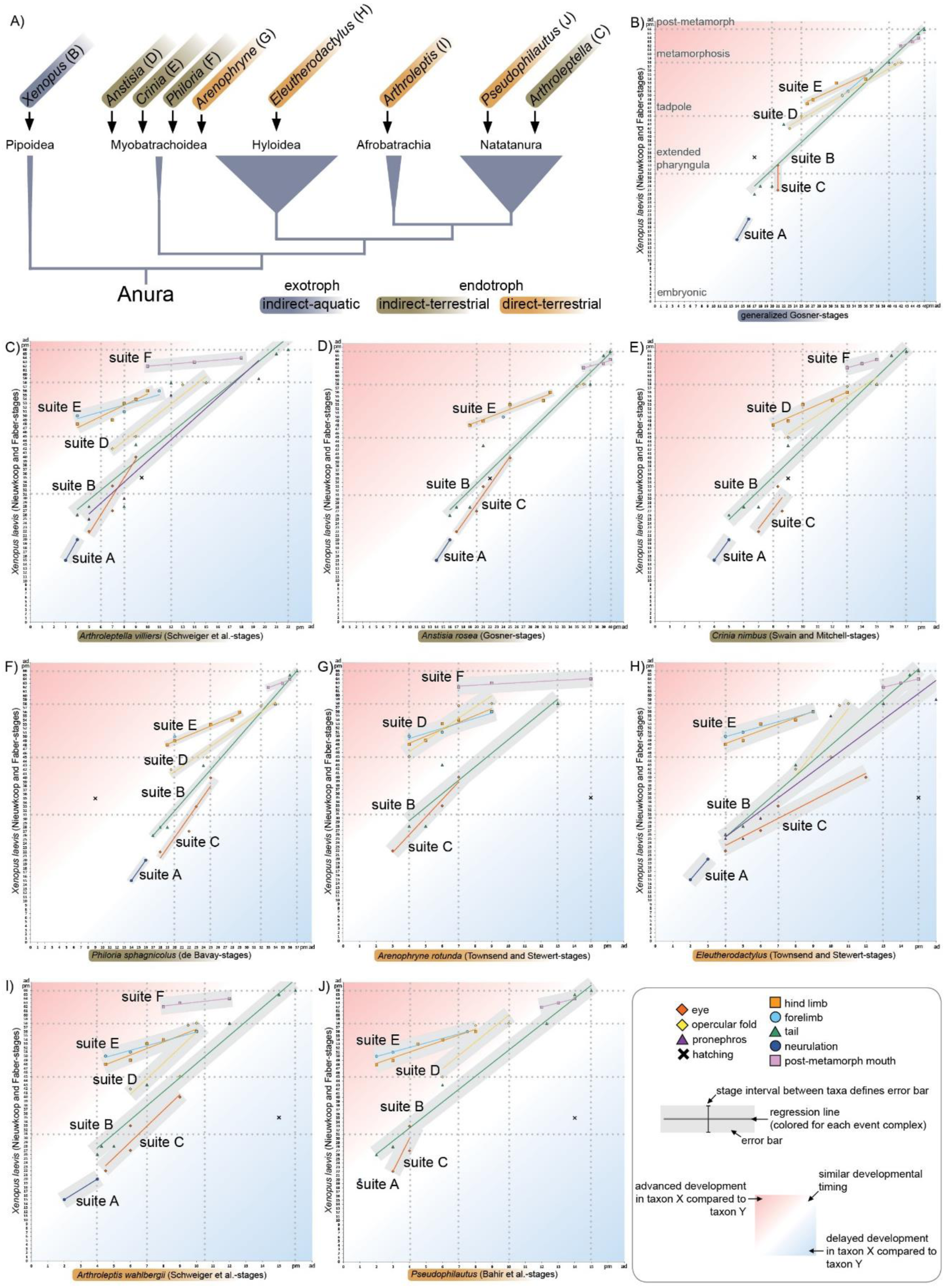
Heterochrony plots of anuran taxa with different reproductive modes. A) Simplified phylogenetic distribution of investigated taxa showing references to the heterochrony plots B) to J).

**Table 4.**
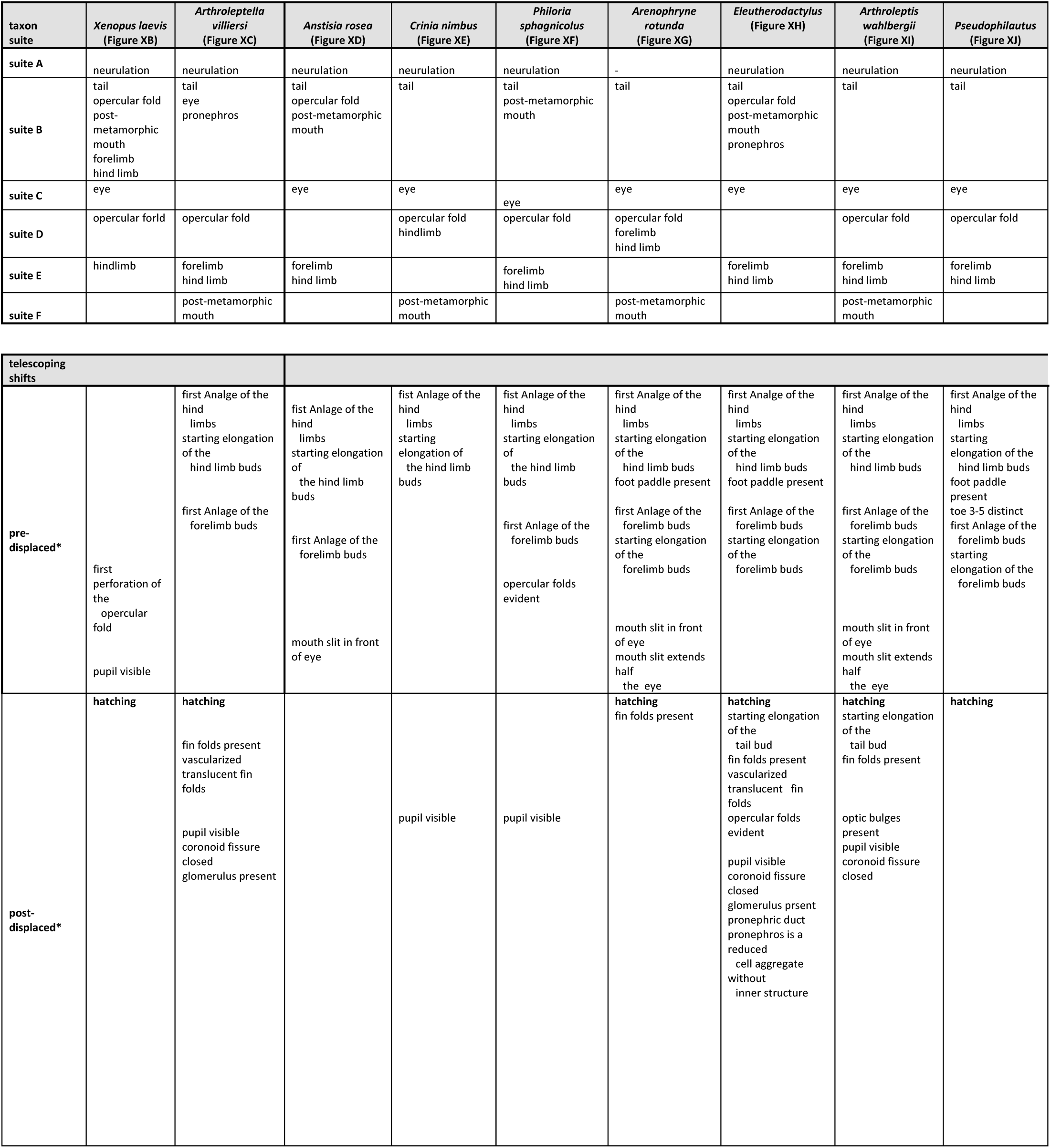
Distribution of developmental events to different isochronic suites deduced from the heterochrony plots and summary of pre-and post-displaced events for every species. *pre-displacement and post-displacement are compared to homologous developmental events in *X. laevis*. In *X. laevis*, it is compared to the generalized Gosner stages.

#### Endotrophic indirect development

The heterochrony plot of *A. villiersi* against *X. laevis* (Figure 3C) reveals a dissociation of the opercular fold developmental suite as isochronic suite D, hind-and forelimb developmental suites together as isochronic suite E and the post-metamorphic mouth developmental suite as isochronic suite F (Table 4). These suites are pre-displaced in *A. villiersi* compared to *X. laevis*. Events that show a clear telescoping shift to an earlier developmental point are the first Anlage of the hind- and forelimbs, the starting elongation of the hindlimb buds and the position of the mouth slit in front of the eye (Table 4). A telescoping shift to a later developmental time point is shown for the presence of fin folds, the vascularization of the translucent fin fold, the visibility of a pupil, closure of the choroid fissure of the eye, the presence of a pronephric glomerulus and hatching (Table 4). The plots of endotrophic indirect developing myobatrachid species reveal a pattern comparable to that of *A. villiersi* (Figure 3D-F). In *A. rosea* and *P. sphagnicolus*, time points for only one developmental event of the forelimb suite are available. Based on the data available, fore- and hind limb development appear co-dissociated (isochoric suite E). For *C. nimbus*, no data on forelimb development are available. In this species, error bars of the hind limb and opercular fold developmental suites cross each other for large parts forming an isochronic suite D. This is different to *P. sphagnicolus*, in which opercular development appears as a separate suite D. Post-metamorphic mouth development only appears as a separate suite F in *C. nimbus*. In contrast to these pre-displaced suites, the eye development suite is slightly post-displaced compared to the other developmental suites appearing as a separate suite C in all three endotrophic indirect developing myobatrachids. A list of the events that exhibit clear telescoping shifts is shown in Table 4. As in *A. villiersi*, early hind limb developmental events are pre-displaced in all three species. Early forelimb development appears pre-displaced in *A. rosea* and *P. sphagnicolus*. In *A. rosea*, the position of the mouth slit in front of the eye is pre-displaced as in *A. villiersi*. In *P. sphagnicolus*, the emergence of the opercular fold is pre-displaced. The only event that shows a clear post-displacement is the visibility of a pupil in *A. nimbus* and *P. sphagnicolus*.

#### Endotrophic direct development

The heterochrony plots of the direct developing *A. rotunda, Eleutherodactylus, A. wahlbergii* and *Pseudophilautus* (Figure 3G-J) show heterochronic patterns that somehow extend the observations from endotrophic indirect species. In all four direct developing species, early events of fore- and hind limb development co-dissociate as an isochronic suite D to an earlier developmental time point (Table 4). In *A. rotunda*, opercular development co-dissociates together with fore- and hindlimb development in an isochronic suite D. This is different to *A. wahlbergii* and *Pseudophilautus*, where it forms a suite D separated from the isochronic suite E including fore- and hindlimb development and to *Eleutherodactylus* where it falls into a suite B together with tail, pronephros and post-metamorphic mouth developmental events (Table 4). In *A. rotunda* and *A. wahlbergii*, events of post-metamorphic mouth development dissociate from other events to an earlier developmental time point, forming a suite F. Eye development is post-displaced in all four examined taxa forming a separate suite C. Events exhibiting a clear telescoping shift are shown in Table 4 for all four taxa. Pre-displacement can be seen for earlier and later events of the fore- and hind limb developmental suites in all four taxa and for events of the post-metamorphic mouth suite in *A. rotunda* and *A. wahlbergii*. Hatching is clearly post-displaced in all four direct developing taxa. Additional post-displaced events include the presence of a fin fold in *A. rotunda, Eleutherodactylus* and *A. wahlbergii*, the starting elongation of the tail bud in *Eleutherodactylus* and *A. wahlbergii* as well as other events of the tail developmental suite in *A. wahlbergii* only. Further post-displaced events include events of the eye developmental suite in *Eleutherodactylus* and *A. wahlbergii* and events of the pronephros development suite in *Eleutherodactylus* (Table 4).

## Discussion

Endotrophic indirect development with a nidicolous tadpole is defined by the presence of a free-living but non-feeding larva hatching from an aquatic or terrestrial egg. The entire larval development and metamorphosis of these nidicolous tadpoles depends solely on yolk reserves deposited in the egg (Thibaudeau & Altig 1999, Altig & Johnston 1989). In some cases, such nidicolous tadpoles are classified as “direct developing” in a broader sense, based on their terrestriality, a certain degree of reduction of typical larval features and the presence of endotrophy (Westrick et al. 2022, Channing et al. 1994, Morgan et al. 1989). Direct development, in a strict sense, is defined by the loss of a free-living larval phase resulting in the emergence of a post-metamorphic juvenile from an often terrestrially deposited egg (Hall and Wake, 1999). Therefore, the time point of hatching (pre-metamorphic in terrestrial indirect developing species and post-metamorphic in terrestrial direct developing species) is the main criterion to distinguish these two reproductive modes from an ecological view (Hall and Wake, 1999). However, this distinction becomes ambiguous when these reproductive modes are compared from a morphological perspective.

The terrestrial endotrophic larva of *A. villiersi* shares external similarities with exotrophic, aquatic larvae e.g. in having a long muscularized tail with a fin, a lateral line system with neuromasts and an opercular fold completely covering the forelimbs (De Villiers, 1929; Morgan et al., 1989; this study). The size, extension and vascularization of the tail fin varies considerably among anurans with terrestrialized reproductive modes. In contrast to the direct-developing *Eleutherodactylus* (Townsend & Stewart, 1985)*, Pseudophilautus* (Bahir et al. 2005) and *Arthroleptis* (Schweiger et al., 2017) showing a weakly muscularized tail with an extended fin fold. A long more or less muscularized tail with a low fin has also been described in the direct-developing myobatrachids *Myobatrachus gouldii* and *Arenophryne rotunda* (Anstis et al., 2007). In the direct-developing bufonid *Osornophryne occidentalis* (Romero-Carvajal et al. 2023), the tail is somewhat similar to *A. villiersi* although it is less muscularized. Some direct-developing taxa share a very expanded fin (Bahir et al. 2005; Anstis et al. 2007, Goldberg et al. 2012; Goldberg and Vera Candioti 2015) whereas others have a lower fin with less vascularization (Townsend and Stewart 1985, Anstis et al. 2011, Schweiger et al. 2017). In the direct-developing strabomantid *Oreobates bardemenos* it has even been shown that there is some phenotypic plasticity with respect to tail fin extension (Salicaet al. 2023). Terrestrial indirect developing species in contrast exhibit a more uniform condition with a muscularized tail and a normal to weakly developed tail fin showing no excess vascularization (Anstis, 2010; de Bavay, 1993; Morgan et al. 1989; Swain & Mitchell, 1996). This pattern of tail muscularization and fin vascularization indicates different functional constraints underlying the evolution of endotrophic indirect and direct developing species. In endotrophic indirect anurans, the tail is still used for moving in the nest requiring some degree of muscularization. It can be assumed that an extended fin would be disadvantageous moving in a very viscous jelly environment of the nest. In endotrophic direct developing anurans in contrast in which the complete metamorphic development happens *in-ovo*, the tail has lost its locomotory function and has gained an increased role in respiration. This resulted in decreased muscle mass and an a strongly vascularized large tail fin to increase to respiratory contact surface with the egg membranes.

An opercular fold completely covering the developing forelimbs has been described in direct-developing myobatrachids (Anstis et al. 2007; Anstis 2008), as well as *Arthroleptis* (Schweiger et al. 2017), *Pseudophilatus* (Bahir et al. 2005) and *O. occidentalis* (Romero-Carvajal et al. 2023). In the direct-developing *Eleutherodactylus coqui* (Callery & Elinson, 2000), *Oreobates barituensis* (Goldberg et al. 2012), *Haddadus binotatus* (Goldberg and Vera Candioti 2015) and *Ischnocnema henselii* (Goldberg et al. 2020) an opercular fold appears only in a reduced state, never covering the forelimbs completely. In exotrophic and some aquatic endotrophic tadpoles, the opercular fold forms a spiracle that is involved in directing the excurrent respiratory water flow and covering the developing forelimbs and thereby creating a more stream-lined body form (Altig & McDiarmid, 1999). The variable differentiation of the opercular fold and its varying heterochronic dissociation from fore limb development indicates a loss of function, first in directing the respiratory water flow and second in covering the forelimbs for a more stream-lined body form, during the evolution of terrestrial endotrophic reproductive modes.

It has been noted previously that fore- and hind limb buds appear earlier and more synchronized in direct-developing frogs compared to aquatic indirect species (Anstis et al., 2011; Elinson, 1994; Schweiger et al., 2017). The heterochrony plots of the investigated direct-developing taxa support this observed pre-displacement and the co-dissociation of early aspects of fore- and hind limb development compared to e.g., tail and eye development. The timing of later events of fore- and hind limb development appear also synchronized but more conserved, showing no pre-displacement (see Table 4). In endotrophic indirect developing taxa, early hind limb development is also pre-displaced (Anstis, 2010; De Villiers, 1929; Swain & Mitchell, 1996). However, detailed data on forelimb development are only available for *A. villiersi*. In this species, the timing of developmental events is similar to direct developing taxa, showing a pre-displaced synchronous timing of early events of fore- and hind limb development. The timing of later events is more conserved (see Table 4). This indicates that limb development in frogs with terrestrial reproductive modes is shaped by functional rather than developmental constraints. While the timing of early events can be shifted profoundly, indicating that there are none or only weak developmental constraints. However, both fore- and hindlimb have to develop coordinated (synchronously). Later developmental events have to become integrated again with other developing aspects of the adult frog body plan resulting in a conserved timing. We interpret this pattern as presence of functional constraints ensuring the proper establishment of the highly specialized adult jumping apparatus.

Another characteristic observed in endotrophic tadpoles is the reduction of tadpole-typical mouth parts. Aquatic endotrophic tadpoles of some cycloramphids occur on rock surfaces near waterfalls and develop keratinized jaws (in addition to other exotrophic tadpole-typical features such as external gills, an opercular fold and a muscularized tail with low tail fins (Colaço and Da Silva 2022)). Similar characteristics are shown in endotrophic tadpoles of the bufonid *Frostius pernambucensis* inhabiting phyototelmata, and the cycloramphid *Eupsophus emiliopugini* although some elements of the oral disc and the buccopharyngeal cavity are reduced (Dubeux et al. 2024). These tadpoles are aquatic, occuring in small water-filled crevices like phytotelmata (*F. pernambucensis,* Dubeux et al. 2024) or in small chambers near streams (*E. emiliopugini,* Vera Candioti et al 2011). In endotrophic tadpoles with complete terrestrial development such as *A. villiersi* (this study), *A. rosea* and other species of e.g., Bufonidae and Hemiphractidae, an oral disc with keratinized jaw structures and external gills are often reduced or even completely lost (Vera Candioti et al. 2011). A complete reduction of tadpole-typical oral structures can be observed in terrestrial direct developing species (Townsend & Stewart 1989, Bahir et al. 2005; Anstis et al. 2007, Goldberg et al. 2012, Schweiger et al 2017 etc.). This reduction of tadpole-typical mouth parts can sometimes result in the pre-displacement of adult mouth formation as seen in *A. villiersi*, *Arthroleptis* and *Arenophryne rotunda* (this study). We also observed the post-displacement of events of early eye development in the endotrophic indirect developing *C. nimbus*, *P. sphagnicolus* and *A. villiersi* as well as the direct developing *Eleutherodactylus* and *Arthroleptis*. Additionally, post-displacement of early tail developmental events are observed in *A. villiersi*, *Arthroleptis* and *Eleutherodactylus*. These results seem to mirror a later importance of the functionality of these structures compared to free-living tadpoles. Interestingly we also observed the pre-displaced of glomerulus development in *A. villiersi* and of glomerus and other pronephric developmental events in *Eleutherodactylus*. This could indicate a differences in the excretory cycle connected to the use of yolk as the only nutritional source during embryonic and larval development. However, more comparative data of the development of the pronephric and excretory system from other exotrophic and endotrophic anurans are needed to test this assumption.

Anurans with an endotrophic reproductive mode show a mosaic of reductions of tadpole-typical features, adaptations to the development within a terrestrial environment (e.g, a nest or the egg) and heterochronic shifts. Some of these heterochronic shifts might be consequences of functional and developmental constraints that underlie the establishment of the adult body plan (e.g., early development of limb buds and conserved developmental timing of later limb morphogenesis). Others seem to be more neutral resulting from a decreased importance of their biological role during certain life phases (e.g., the post-displaced development of the eye). Interestingly, while the degree of reduction of tadpole-typical features seems to correlate with the degree of terrestriality of development, patterns of heterochronic shifts appear more complex. *Arthroleptella villiersi* for example, classified as endotrophic indirect developing, shows a pattern of post-displaced events very similar to the direct developing *Eleutherodactylus* and *Arthroleptis*. On the contrary, the direct developing *A. rotunda* and *P. silus* show less post-displaced events and thereby a heterochronic pattern more similar to indirect developing species. The data further indicate that the categories endotrophic indirect and direct development are based on a continuum/spectrum of reproductive modes that show specific profiles of developmental repatterning. More detailed data of completely endotrophic and, even more interesting, partially endotrophic species can help to further explorer this spectrum and increase our understanding of the heterochrony as a pattern as well as a process underlying anuran reproductive diversity.

## Acknowledgements

We thank Alan Channing and Andrew Turner for their advice, help in the field, and hospitality. Permits to collect were issued by CapeNature (permits No. 0035-AAA004-01041, 0056-AAA041-00073). This research was made possible through a German Science Foundation (DFG) grant to HM (MU 2914/2-1) and financial assistance by the Friedrich Schiller University to HM, which is gratefully acknowledged.

## References

Altig, R., Johnston, G.F. (1989). Guilds of anuran larvae: Relationships among developmental modes, morphologies, and habitats. Herpetological Monographs 3: 81–109.

Anstis, M. (2008). Direct development in the Australian myobatrachid frog *Metacrinia nichollsi* from Western Australia. Records of the Western Australian Museum, 24: 133–150.

Anstis, M. (2010). A comparative study of divergent embryonic and larval development in the Australian frog genus *Geocrinia* (Anura: Myobatrachidae). Records of the Western Australian Museum, 25(4): 399–440.

Anstis, M., Parker, F., Hawkes, T., Morris, I., & Richards, S. J. (2011). Direct development in some Australopapuan microhylid frogs of the genera *Austrochaperina*, *Cophixalus* and *Oreophryne* (Anura: Microhylidae) from northern Australia and Papua New Guinea. Zootaxa, 3052: 1–50.

Anstis, M., Roberts, J.D., Altig, R. (2007). Direct development in two Myobatrachid Frogs, *Arenophryne rotunda* Tyler and *Myobatrachus gouldii* Gray, from Western Australia. Records of the Western Australian Museum 23: 259–271.

Bahir, M. M., Meegaskumbura, M., Manamendra-Arachchi, K., Schneider, C. J., & Pethiyagoda, R. (2005). Reproduction and terrrestrial direct development in Sri Lankan Shrub Frogs (Ranidae: Rhacophorinae: Philautus). Raffles Bulletin of Zoology, 12: 339–350.

Bittencourt-Silva, G.B., Conradie, W., Siu-Ting, K., Tolley, K.A., Channing, A., Cunningham, M., Farooq, H.M., Menegon, M., Loader, S.P. (2016) The phylogenetic position and the diversity of the enigmatic mongrel frog *Nothophryne* Poynton 1963 (Amphibian: Anura). Molecular Phylogenetics and Evolution 99: 89–102.

Brink, H.E. (1939). A histological and cytological investigation of the thyroids of *Arthroleptella bicolor villiersi* and *Bufo angusticeps* during normal and accelerated metamorphosis. Proceedings of the Linnean Society of London 151: 120–125.

Brink, H.t., Onstein, R.E., de Roos, A.M. (2020). Habitat deterioration promotes the evolution of direct development in metamorphosing species. Evolution 74 (8):1826–1850.

Caldwell, J.P., Lima, A. P. (2003). A new amazonian species of *Colostethus* (Anura: Dendrobatidae) with a nidicolous tadpole. Herpetologica 59 (2): 219–234.

Callery, E.M., Fang, H., Elinson, R.P. (2001). Frogs without polliwogs: evolution of anuran direct development. BioEssays 23: 233–241.

Callery, E. M., & Elinson, R. P. (2000). Opercular development and ontogenetic reorganization in a direct-developing frog. Development Genes and Evolution, 210: 377–381.

Channing, A. (2004a). The genus Arthroleptella Hewitt, 1926 (Family Petropedetidae), p. 206–207. In: Atlas and Red Data Book of the Frogs of South Africa, Lesotho and Swasiland. L. R. Minter, M. Burger, J. A. Harrison, H. H. Braack, P. J. Bishop, and D. Kloepfer (eds.). SI/MAB Series #9. Smithsonian Institution, Washington, D.C.

Channing, A. (2004b). *Arthroleptella* villiersi Hewitt, 1935, p. 218–219. In: Atlas and Red Data Book of the Frogs of South Africa, Lesotho and Swasiland. L. R. Minter, M. Burger, J. A. Harrison, H. H. Braack, P. J. Bishop, and D. Kloepfer (eds.). SI/MAB Series #9. Smithsonian Institution, Washington, D.C.

Channing, A., Hendricks, D., Dawood, A. (1994). Description of a new moss frog from the south-western Cape (Anura: Ranidae: *Arthroleptella*). South African Journal of Zoology 29 (4): 240–243.

Chuliver, M., Agnolín, F. L., Scanferla, A., Aranciaga Rolando, M., Ezcurra, M. D., Novas, F. E. & Xu, X. (2024). The oldest tadpole reveals evolutionary stability of the anuran life cycle. Nature 636: 138–142.

Colaço, G., da Silva, H. R. (2022). Finding a pathway through the rocks: the role of development on the evolution of quasi-terrestriality and the origin of endotrophism in cycloramphids (Anura), Biological Journal of the Linnean Society 137 (2): 294–323.

Crump, M. L. (2015) Anuran Reproductive Modes: Evolving Perspectives. Journal of Herpetology 49(1): 1–16.

de Bavay, J. M. (1993). The developmental stages of the Sphagnum frog, *Kyarranus sphagnicolus* Moore (Anura, Myobatrachidae). Australian Journal of Zoology, 41(2): 151–201.

de Villiers, C.G.S. (1929). The development of a species of *Arthroleptella* from Jonkershoek, Stellenbosch. South African Journal of Science 26: 481–510.

du Preez, L., Carruthers, V. (2017) Frogs of Southern Africa, a complete Guide. Struik Nature.

Dubeux, M.J.M., do Nascimento, F.A.C. & Dias, P.H. (2024). Larval morphology of *Frostius pernambucensis* (Anura): contribution of larval characters for the systematics of the family Bufonidae and evolution of endotrophic tadpoles. Zoomorphology 143: 159–187.

Duellman, W. E., & Trueb, L. (1994). Biology of amphibians: JHU press.

Goldberg, J.,& Vera Candioti, F. (2015). A tale of a tail: variation during the early ontogeny of *Haddadus binotatus* (Brachycephaloidea: Craugastoridae) as compared with other direct developers. Journal of Herpetology, 49: 479–484.

Goldberg, J., Vera Candioti, F., & Akmentins, M. S. (2012). Direct developing frogs: ontogeny of *Oreobates barituensis* (Anura: Terrarana) and the development of a novel trait. Amphibia-Reptilia, 33: 239–250.

Goldberg, J., Taucce, P. P. G., Quinzio, S. I., Haddad, C. F. B., & Vera Candioti, F. (2020). Increasing our knowledge on direct developing frogs: The ontogeny of *Ischnocnema henselii* (Anura: Brachycephalidae. Zoologischer Anzeiger, 284: 78–87.

Gosner, K.L. (1960). A simplified table for staging anuran embryos and larvae with notes on identification. Herpetologia 16: 183–190.

Grosso, J., Baldo, D., Cardozo, D., Kolenc, F., Borteiro, C., de Oliveira, M. I., … & Vera Candioti, F. (2019). Early ontogeny and sequence heterochronies in Leiuperinae frogs (Anura: Leptodactylidae). PLoS One, 14(6), e0218733.

Elinson, R. P. (2009). Nutritional endoderm: a way to breach the holoblastic–meroblastic barrier in tetrapods. Journal of Experimental Zoology Part B: Molecular and Developmental Evolution, 312(6): 526–532.

Haddad C, Prado CPA. 2005 Reproductive modes in frogs and their unexpected diversity in the Atlantic Forest of Brazil. BioScience 55: 207–217. (doi: 10.1641/0006-3568(2005)055[0207:RMIFAT]2.0.CO;2)

Hanken, J. (2003): Direct development. In: Keywords and Concepts in Evolutionary Developmental Biology. Edited by B.K. Hall and W.M Olson. Cambridge, Harvard University Press, pp. 97–102

Hanken, J. (2006). Direct development. In B. K. Hall & W. M. Olson (Eds.), Keywords and concepts in evolutionary developmental biology (pp. 97–102): Harvard University Press.

Hanken, J., Klymkowsky, M.W., Summers, C.H., Seufert, D.W., Ingebrigtsen, N. (1992). Cranial ontogeny in the direct-developing Frog, *Eleutherodactylus coqui* (Anura: Leptodactylidae), analyzed using Whole-Mount Immunohistochemistry. Journal of Morphology 211: 95–118.

IUCN SSC Amphibian Specialist Group (2013). Arthroleptella villiersi. The IUCN Red List of Threatened Species 2013: e.T58063A18403581. 10.2305/IUCN.UK.2013-2.RLTS.T58063A18403581.en. Accessed on 23 January 2025.

Juncá, F.A., Altig, R., Gascon, C. (1994). Breeding Biology of *Colostethus stepheni*, a Dendrobatid Frog with a Nontransported Nidicolous Tadpole. Copeia, Vol. 1994, No. 3: 747–750.

Lutz, B. (1947). Trends towards Non-Aquatic and Direct Development in Frogs. Copeia 4: 242–252.

Morgan, B. E., Passmore, N.I., Fabian, B.C. (1989). Metamorphosis in the frog *Arthroleptella lightfooti* (Anura, Ranidae) with emphasis on neuro-endocrine mechanism. In: Bruton, M.N. (ed.), Alternative Life-History Styles of Animals: 347–370. Kluwer Academic Publishers, Dordrecht.

Müller, H., Liedtke, H. C., Menegon, M., Beck, J., Ballesteros-Mejia, L., Nagel, P., & Loader, S. P. (2013). Forests as promoters of terrestrial life-history strategies in East African amphibians. Biology Letters, 9(3), 20121146.

Raff, R.A., Wray, G.A. (1989). Heterochrony. Developmental mechanisms and evolutionary results. Journal of Evolutionary Biology 2: 409–434.

Romero-Carvajal A, Negrete L, Salazar-Nicholls MJ, Vizuete K, Debut A, Dias PH, Vera Candioti F (2023). Direct development or endotrophic tadpole? Morphological aspects of the early ontogeny of the plump toad *Osornophryne occidentalis* (Anura: Bufonidae). J Morphol. 284(5).

Salica, M. J., Goldberg, J., Akmentins, M. S., & Candioti, F. V. (2023). Exceptional features of the embryonic ontogeny of a direct-developing Robber frog. Journal of Zoology, 320(2): 120–130.

Schlosser, G. (2001). Using heterochrony plots to detect the dissociated coevolution of characters. Journal of Experimental Zoology, 291(3): 282–304.

Schlosser, G., & Roth, G. (1997). Evolution of nerve development in frogs; pp. 94–112. Brain, Behavior and Evolution, 50(2): 94-112.

Schweiger, S., Naumann, B., Larson, J. G., Möckel, L. & Müller, H. (2017). Direct development in African squeaker frogs (Anura: Arthroleptidae: *Arthroleptis*) reveals a mosaic of derived and plesiomorphic characters. Organisms, Diversity and Evolution 17: 693–707.

Swain, R., & Mitchell, N. (1996). Terrestrial development in the Tasmanian frog, *Bryobatrachus nimbus* (Anura: Myobatrachinae): larval development and a field staging table. Papers and Proceedings of the Royal Society of Tasmania, Volume 130: 1.

Thibaudeau, G., Altig, R. (1999). Endotrophic anurans. Development and Evolution. In: Tadpoles – the biology of anuran larvae. Edited by R. W. Mc Diarmid and R. Altig. Chicago University Press, USA, pp. 24–51.

Townsend, D. S., & Stewart, M. M. (1985). Direct development in *Eleutherodactylus coqui* (Anura: Leptodactylidae): a staging table. Copeia 1985: 423–436.

Turner, A., Channing, A. (2017) Three new species of *Arthroleptella* Hewitt, 1926 (Anura: Pyxicephalidae) from the Cape Fold Mountains, South Africa. African Journal of Herpetology 66: 53–78.

van der Meijden, A., Crottini, A., Tarrant, J., Turner, A., Vences, M. (2011). Multi-locus phylogeny and evolution of reproductive modes in the Pyxicephalidae, an African endemic clade of frogs. African Journal of Herpetology 60: 1–12.

Vera Candioti, M.F., Nuñez, J.J., Úbeda, C. (2011). Development of the nidicolous tadpoles of *Eupsophus emiliopugini* (Anura: Cycloramphidae) until metamorphosis, with comments on systematic relationships of the species and its endotrophic developmental mode. Acta Zoologica (Stockholm) 92: 27–45.

Westrick, S.E., Laslo, M., Fischer, E.K. (2022). The Natural History of Model Organisms: The big potential of the small frog *Eleutherodactylus coqui*. eLife: 1–20.

